# Accuracy of computed tomography in diagnosing temporomandibular joint osteoarthritis relative to histopathological findings – an ex–vivo study of 41 horses

**DOI:** 10.1101/2024.11.01.621628

**Authors:** Tomasz Jasinski, Łukasz Zdrojkowski, Bernard Turek, Michał Kaczorowski, Bartosz Pawliński, Walter Brehm, Małgorzata Domino

## Abstract

Computed tomography (CT) is used to support the diagnosis of equine temporomandibular joint (TMJ) disease; however, its diagnostic accuracy remains unclear. This study aimed to evaluate the relationship between CT findings and histopathological manifestations of osteoarthritis (OA) in equine TMJs. 82 TMJs were CT–imaged, sampled, grouped into age–related and OA–related groups, and analyzed for frequency distributions, correlations, and CT–based TMJ OA diagnosis. CT findings were observed in 79% of joints, including ‘CT anatomical variations’ considering to reflect age–related remodeling. Only 50% of joints showed co–occurrence of CT findings and histopathological manifestations of OA, confirming that not all CT findings are indicative of disease. Including all CT findings in the CT–based diagnosis of TMJ OA yielded a specificity of 0.41 (95% CI: 0.26–0.58), suggesting a high rate of false–positive diagnoses. Excluding all ‘CT anatomical variations’ resulted in a sensitivity of 0.56 (95% CI: 0.40–0.72), indicating a substantial number of false–negative diagnoses. However, inclusion of specific ‘CT anatomical variation’ – subchondral bone cysts – into the studied CT–based diagnosis increased sensitivity to 0.79 (95% CI: 0.62 to 0.89) while maintaining high specificity of 0.92 (95% CI: 0.80–0.98). Including this subset of CT findings in the diagnosis of equine TMJ OA may improve the accuracy of disease detection; however, the clinical relevance of the present cadaver investigation needs to be confirmed in in vivo studies.

## 1 INTRODUCTION

Diseases of the temporomandibular joint (TMJ) in horses have been suggested to be rare or rarely reported [1,2]. Their prevalence in the equine population is unknown; however, clinical recognition is increasing [3]. The TMJ facilitates movement of the mandibular condyle relative to the squamous part of the temporal bone within a deep mandibular fossa [4]; therefore, disease–related dysfunctions may lead to various clinical symptoms, ranging in severity from mild behavioral issues [5–7] to moderate masticatory difficulties [8,9] and severe inability to open [10,11] or close [12,13] the mouth.

Among TMJ diseases, osteoarthritis (OA) may develop as a primary condition [5–7,11,14,15] or secondarily to fractures [16] or septic arthritis [1,10,17]. OA involves the progressive degradation of the fibrocartilage covering the articular surfaces of the TMJ, ultimately leading to changes in the underlying subchondral bone. Increasingly, horses presented to veterinarians due to milder symptoms – such as poor performance [6,7], aversive behaviors [5,6], head–shaking syndrome [5,18], and resistance or reluctance to work on the bit [6,7] – are suspected of having TMJ OA. Consequently, TMJ OA is discussed more frequently [2,19] as a potential cause of nonspecific clinical symptoms associated with decreased performance in equine athletes [2,5–7,11,14,15]. However, the initial diagnosis of primary TMJ OA – which should include detailed history, clinical symptoms, dental examination, and exclusion of other more common potential reasons of symptoms [2] – requires confirmation using diagnostic imaging modalities [5– 7,11,14–16,20] and, when possible, intra–articular analgesia [6,7].

Among diagnostic imaging modalities, ultrasonography [5,11], conventional radiography [5,11,16], and/or computed tomography (CT) [5–7,14,15,20] are used clinically for the diagnosis of TMJ OA in horses. Given the anatomical complexity and significant superimposition of anatomical structures in the TMJ region, image clarity of ultrasonography [21] and conventional radiography is limited [22], even when specific TMJ radiographic views are employed [23–25]. Therefore, the advantages of CT over conventional radiography are emphasized [1,10,16,17,22]. Moreover, with recent advances in performing CT imaging of the equine head under standing sedation [26,27], CT examinations in horses are expected to increase. Consequently, CT is expected to be widely used to assess TMJ diseases, as it has been in recent research studies [15,18,20] and clinical case reports [5–7,14]. The increased availability and safety of CT imaging also align with the growing number of case reports of primary TMJ OA [2,5–7,11,14,15], which have shown responses to intra–articular medication [2,5–7] and have been associated with head–shaking syndrome [5,15] or more subtle riding problems [2,6,7]. Although almost all of these horses were diagnosed using CT [5–7,14,15], histopathological confirmation of TMJ OA has been reported in only two cases [5,14], and the diagnostic accuracy of CT for equine TMJ OA has not yet been determined [3].

In humans, the accuracy of clinical diagnosis of TMJ OA when unsupported by diagnostic imaging is 0.55 in sensitivity and 0.61 in specificity [28], whereas no similar data have been published for horses [3]. When human TMJ assessment is expanded by CT imaging, a diagnostic sensitivity increases to 0.67–0.90 and specificity to 0.73–0.93 [29]. Moreover, in humans, the Diagnostic Criteria for Temporomandibular Disorders (DC/TMD) are well–defined, indicating that the TMJ OA diagnosis should be confirmed by at least one of the following CT findings: subchondral bone cyst, erosion, generalized subchondral sclerosis, or osteophythosis [28]. Given that in humans flattening of the mandibular condyle and/or cortical sclerosis may represent normal variation, aging, remodeling, or a precursor to frank OA, they are considered indeterminant findings for the disease following DC/TMD guidelines [28].

In horses, similar CT findings – such as flattening of the mandibular condyle, scattered regions of hypodensity, bone cysts, mandibular clefts, medial enthesophytes, and intra–articular disc mineralization – have been reported as incidental and may not be manifested clinically in certain horses [20]. These findings, referred to as ‘CT anatomical variation’ [20], differ from normal radiological anatomy [30], and most are thought to reflect age–related remodeling of the TMJ in horses, including those without moderate to severe clinical symptoms of TMJ dysfunction [20]. Therefore, inclusion of such findings in CT–based assessment may affect the diagnostic accuracy for TMJ OA. However, recent case reports have linked certain ‘CT anatomical variations’ – such as subchondral bone cysts in the mandibular condyle [6,7] and mineralization of the intra–articular disc [5,6] – with mild performance–related clinical symptoms of TMJ diseases, thereby increasing interest in the radiological manifestations of TMJ OA and renewing discussion on their clinical significance [2,19]. Notably, the main difference in CT manifestation of TMJ OA listed in humans [28] and horses [20] refers to subchondral bone cysts, which in humans are considered CT–positive for OA [28], while in horses were initially suggested as ‘CT anatomical variations’ with presently unknown clinical significance [20].

To address the gap in understanding the contribution of specific CT findings to the diagnosis of equine TMJ OA, the study aims to describe the relationship between CT findings in equine TMJs and histopathological manifestations of OA, focusing on their frequency distributions and correlations with horses’ age. Subsequently, this study aims to evaluate the accuracy of CT–based diagnosis of TMJ OA, considering three subsets of CT findings – one including all CT findings, second including CT findings without equine ‘CT anatomical variations’ [20], and third including CT findings with bone cysts [28] but without remaining ‘CT anatomical variations’ [20].

## 2 METHODS

### 2.1 Selection and description of subjects

This prospective cross–sectional study was conducted on 50 equine cadaver heads collected from horses at a commercial slaughterhouse in Rawicz, Poland. The study was carried out between May and June 2023. At the time of slaughter using a physical method (penetrating captive bolt), the age of each horse was recorded. The horse heads (mean ± standard deviation (SD) length 52.5 ± 3.2 cm, width 21.3 ± 1.3 cm, and height 30.8 ± 2.0 cm) were collected post mortem, which does not fall under legislation governing for protection of animals used for scientific purposes, according to national law (Dz. U. 2015 poz. 266) and EU Directive (2010–63–EU). Therefore, ethical committee approval was not required. The cadaver heads underwent TMJ palpation, CT examination, and tissue sample collection for histopathological assessment. The interval between slaughter and completion of the research protocol for each head did not exceed 4 h. The results of TMJ palpation, CT examination, and histopathological assessment were used to determine joint inclusion or exclusion and for data grouping. The eligibility criteria required the presence of a macroscopically intact joint capsule, a complete CT scan of the TMJ, and complete histopathological slides without signs of autolysis. The exclusion criteria included the presence of a wound in the TMJ region, disruption of the joint capsule, fracture of the coronoid process, mandibular condyle or mandibular fossa, TMJ luxation, the presence of dentigerous cysts in the TMJ region, radiological signs of TMJ neoplasia, and missing or poor– quality tissue samples on histopathology slides.

### 2.2 CT image collection and analysis

CT examination was performed using a multi–slice scanner (Revolution CT, 64–rows, GE Healthcare, Chicago, IL, USA). The cadaver heads were scanned in ventral recumbency with close months using the following acquisition parameters: helical scan mode; current 275 mA; voltage range 70–140 kV in GSI–QC mode; gantry rotation 0.08 s (HE+); table travel 39.4 mm/rotation; pitch 0.984:1; and slice thickness 2.5 mm. The scan length and duration were adjusted to the size of the head, with the scan length covering the area from the rostral aspect of the lips to the caudal aspect of the occipital bone, and the scan duration not exceeding 20 s. Thus, the number of slices was tailored to the size of each head. The CT scans were processed using the AW workstation and VolumeShare 7 software (GE Healthcare, Chicago, IL, USA) to generated detailed reconstruction with a monovoltage of 70 kV and a slice thickness of 0.625 mm. The reconstructed images were saved in DICOM format and reviewed using Osirix MD software version 12.0 (Pixmeo SARL, Bernex, Switzerland) and multiplanar reconstruction in both bone window (width 1500, level 300) and soft tissue window (width 300, level 50). The images were independently, randomly, and simultaneously reviewed by two Polish board–certified veterinary diagnostic imaging (PCVDI) specialists (T.J. and M.D.). Both observers were blinded to the horse data and histopathological findings. Any disagreements were resolved by a Polish board–certified equine disease (PCED) specialist (B.T.), and the consensus assessment was used as the final result.

To standardize the CT image review, a CT findings extraction template was developed based on previously published studies [5–7,14,15,20]. The template is presented in Table 1. The CT findings for each joint was annotated as present (1) or absent (0), including changes in the following categories: bone surface (flattening; irregularity), subchondral bone (scattered, spherical or linear regions of hypodensity; with or without surrounding hyperdensity (sclerosis); general hyperdensity (sclerosis) within subchondral bone), bone margins (enthesophytes; marginal osteophytes), and joint space (narrowing; widening; hyperdensity). The localization of specific CT findings within the mandibular condyle, mandibular fossa, and joint space (including the intra–articular disc and periarticular soft tissues) was also recorded. If no CT findings were reported, the joint was annotated as normal in a given category.

**Table 1.** Computed tomography (CT) findings observed in equine temporomandibular joints (TMJs). CT findings were localized in the 698 mandibular condyles (MC), mandibular fossa (MF), and joint space (JS). Each CT finding is supported by references in which it has been 699 previously described [5–7,14,15,20]. ‘CT anatomical variations’ are indicated by asterisk.

| Bone surface |  |  | Subchondral bone |  |  | Bone margins |  |  | Joint space |  |  |
| --- | --- | --- | --- | --- | --- | --- | --- | --- | --- | --- | --- |
| 1 | Normal | no flattening and irregularity [7,20] | 1 | Normal | no hypodensity and hyperdensity [7,20] | 1 | Normal | lack of osteophytes and enthesophytes [7,20] | 1 | Normal | no hyperdensity and narrowing / widening [7,20] |
| 2 | Flatt MC* | flattening of MC [14,20] | 2 | ScHD+Scl MC* | scattered regions of hypodensity in MC with surrounding hyperdensity [15,20] | 2 | Medial Ent MC* | enthesophytes on the medial aspect of MC [20] | 2 | Narrow JS | narrowing of the joint space [15] |
| 3 | Irr MC | irregularity of MC [15,20] | 3 | ScHD–Scl MC* | scattered regions of hypodensity in MC without surrounding hyperdensity [20] | 3 | Lateral Ost MC | marginal osteophytes on the lateral aspect of MC | 3 | Wide JS | widening of the joint space [15] |
| 4 | Irr MF | irregularity of MF | 4 | ScHD+Scl MF * | scattered regions of hypodensity in MF with surrounding hyperdensity [15,20] | 4 | Rostral Ost MC | marginal osteophytes on the rostral aspect of MC [14] | 4 | Min Disc* | hyperdensity of the intra-articular disc [5,6,20] |
|  |  |  | 5 | ScHD–Scl MF * | scattered regions of hypodensity in MF without surrounding hyperdensity [20] | 5 | Lateral Ost MF | marginal osteophytes on the lateral aspect of MF [5] | 5 | Osseous JS | hyperdense osseous fragments in the joint space [14,15] |
|  |  |  | 6 | SphHD+Scl MC* (bone cyst) | spherical region of hypodensity in MC with surrounding hyperdensity [6,7,15,20] |  |  |  | 6 | Min Soft | hyperdensity of soft tissues of the periarticular region [20] |
|  |  |  | 7 | SphHD–Scl MC* (bone cyst) | spherical region of hypodensity in MC without surrounding hyperdensity [20] |  |  |  |  |  |  |
|  |  |  | 8 | SphHD+Scl MF * (bone cyst) | spherical region of hypodensity in MF with surrounding hyperdensity [20] |  |  |  |  |  |  |
|  |  |  | 9 | SphHD–Scl MF * (bone cyst) | spherical region of hypodensity in MF without surrounding hyperdensity [20] |  |  |  |  |  |  |
|  |  |  | 10 | Mandibular cleft in MC* | linear region of hypodensity in MC with surrounding hyperdensity [20] |  |  |  |  |  |  |
|  |  |  | 11 | Scl MC | hyperdensity within the subchondral bone [5] |  |  |  |  |  |  |

Footnotes: Cleft = linear region of hypodensity surrounded by sclerosis; Disc = intra-articular disc; Ent = enthesophytes; Flatt = flattening; Irr = irregularity; JS = the joint space; MC = the mandibular condyle; MF = the mandibular fossa in the squamous part of the temporal bone; Min = mineralization (hyperdensity); Ost = osteophytes; ScHD = scattered regions of hypodensity; Scl = sclerosis (hyperdensity); SphHD = spherical region of hypodensity; Soft = soft tissues; + = with; – = without; \* = 'CT anatomical variations'.

Flattening was defined as a reduction or even loos in the globose shape of the curved articular surface of the mandibular condyle, resulting in a more T–shaped appearance [14,20], whereas irregularity was defined as an irregular subchondral bone contour [15,20]. Subchondral bone hypodensity was described as scattered [15,20], spherical [6,7,15,20], or linear [20]. Mandibular clefts were defined as subchondral linear regions of hypodensity surrounded by sclerosis in the mandibular condyle [20]. Scattered and spherical regions of hypodensity were further categorized based on the surrounding hyperdensity (sclerosis), as those with surrounding sclerosis and those with little to no surrounding sclerosis [20]. Spherical regions of hypodensity, suggestive of subchondral cystic lesions (bone cysts), were assessed as open to the articular surface of the bone or close to the articular surface of the bone [20].

Subchondral sclerosis was defined as general radiodensity (hyperdensity) within subchondral bone [5]. Medial enthesophytes were defined as irregularities and modeling of the medial margin of the mandibular condyle at the insertion of the lateral pterygoid muscle [20], whereas osteophytes referred to irregularities of the lateral or rostral margins of the mandibular condyle or mandibular fossa [14,20]. Changes in the joint space were described in terms of size as narrowing [15] or widening [15], and in terms of density as hyperdenity [5,6,14,15,20]. Point–like, linear, or diffuse hyperdensity located within the intra– articular disc was annotated as intra–articular disc mineralization [5,6,20], whereas round or irregular hyperdensity located outside the intra–articular disc was considered suggestive of osseous fragments within the joint space [14,15]. Additionally, any hyperdensity of the periarticular soft tissues [20] was recorded in this category.

Noteworthy, specific CT findings were considered potential ‘CT anatomical variations’ [20] and were marked with asterisks. They included flattening of the mandibular condyle, scattered regions of hypodensity, bone cysts, mandibular clefts, medial enthesophytes, and intra–articular disc mineralization [20].

### 2.3 Tissue samples collection and analysis

From each joint, four tissue samples were collected, representing the articular cartilage of the mandibular condyle, the articular cartilage of the mandibular fossa, the intra–articular disc, and the joint capsule including the synovium. The samples were immersed in 10% neutral buffered formalin (NBF solution; Sigma–Aldrich, Burlington, MA, USA) and fixed for 48 hours at room temperature. Samples were then embedded in paraffin (paraffin wax; Sigma–Aldrich, Burlington, MA, USA), cut into 5 μm sections, and processed for standard histological staining using a hematoxylin–eosin (HE) protocol (hematoxylin 3801520E; Leica, Wetzlar, Germany; eosin, HT1103128; Sigma–Aldrich, Burlington, MA, USA). The HE– stained slides were scanned using a Pannoramic 250 Flash III scanner (3DHistech, Budapest, Hungary) and evaluated with Pannoramic Viewer version 1.15.4 (3DHistech, Budapest, Hungary), at magnification ranging from 40× to 400×. The slides were independently, randomly, and simultaneously reviewed by two observers (Ł.Z. and M.D.), both holding Master’s degree in animal science and trained in histopathological assessment. Both observers were blinded to the horse data and CT findings. Any disagreements were resolved by a Polish board–certified equine disease (PCED) specialist (B.P.), and the consensus assessment was used as the final result.

To standardize the HE–stained slides assessment, histopathological findings for each joint were graded according to the categories described by McIlwraith [31]. These included evaluation of the articular cartilage (chondrocyte necrosis; cluster formation; fibrillation/fissuring; focal cell loss) and joint capsule (cellular infiltration; vascularity; intimal hyperplasia; subintimal edema; subintimal fibrosis) [31]. Additionally, each joint was assessed for alterations in the intra–articular disc, including: cellular infiltration, focal cell loss [32], chondroid metaplasia [5,32], and chondro–osseous metaplasia [5,32]. To facilitate analysis of the relationship between CT and histopathological findings, the Mankin score [31] was adapted to a binary scale, where 0 represented normal and 1 indicated the presence of alterations. Accordingly, histopathological findings for each joint and each category was annotated as either absent (0) or present (1).

### 2.4 Statistical analysis

The statistical analysis was performed using GraphPad Prism version 6 (GraphPad Software Inc., San Diego, CA, USA).

#### 2.4.1 CT and histopathological findings in equine TMJs across age–related groups

TMJs were grouped by horses’ age into three age–related groups: (1) 1–4 years group, (2) 5–15 years group, and (3) >15 years group. The criterion for classification into each groups was the horse’s age. For each age– related group, descriptive statistics were calculated, including the median and range of age, as well as the number and percentage of joints with specific CT and histopathological findings. For each age–related group, the frequency distribution of CT and histopathological findings was calculated separately. The frequency distributions were then compared between age–related groups using a Chi–square test. Differences were considered significant at p < 0.05.

The correlations between the horses’ age and the occurrence of CT findings, as well as the horses’ age and histopathological findings, were calculated separately. As the CT and histopathological findings were represented by the binary data, the Spearman’s rank correlation coefficient (ρ) was used. Correlations were considered significant at p < 0.05. The strength of the correlations was interpreted as weak (≤ 0.40), moderate (0.41–0.70), strong (0.71–0.90), or very strong (≥ 0.91). The direction of the correlation was interpreted as positive (> 0) or negative (< 0) [33].

#### 2.4.2 Relationship between CT and histopathological findings in equine TMJs

The correlations between the CT and histopathological findings were calculated for all TMJs, with no additional grouping. The Spearman’s rank correlation coefficient (ρ) was used. Correlations were considered significant at p < 0.05. The strength of the correlations was interpreted as weak (≤ 0.40), moderate (0.41– 0.70), strong (0.71–0.90), or very strong (≥ 0.91). The direction of the correlation was interpreted as positive (> 0) or negative (< 0) [33].

Subsequently, TMJs were grouped by histopathological findings into a TMJ OA group and OA–free TMJ group. The criteria for classification into the TMJ OA group were both: (1) the presence of at least one histopathological finding listed in the Osteoarthritis Research Society International (OARSI) recommendations [31] and (2) histopathological evidence of active inflammation of the joint capsule (cellular infiltration), allowing differentiation between osteoarthritis and osteoarthrosis [28,34]. Joints were assigned to the OA–free TMJ group when at least one above criterion was not met. For both TMJ OA– related groups, the frequency distribution of CT findings was calculated. The frequency distributions were compared between TMJ OA–related groups using a Chi–square test. Differences were considered significant at p < 0.05.

#### 2.4.3 Accuracy of CT in diagnosing TMJ OA

The accuracy of CT–based recognition of TMJ OA was calculated across three subgroups: subgroup 1 – including all CT findings; subgroup 2 – including CT findings excluding ‘CT anatomical variations’ [20], and subgroup 3 – including CT findings with bone cysts [28] but excluding other ‘CT anatomical variations’ [20]. TMJ was annotated as CT–positive for OA when ad least one CT finding in each subgroup was presented (1). In each case, the group with histopathologically confirmed TMJ OA (TMJ OA group) served as a reference. Sensitivity (Se), specificity (Sp), positive predictive value (PPV), and negative predictive value (NPV) were calculated across a range from 0.1 to 1.0 using standard formulae [35]. The area under curve (AUC), represented the area under the receiver operating characteristic (ROC) curve, was also calculated. Confidence intervals (95% CI) were determined. Accuracy metrics were calculated using diagnostic test (2×2 table) in MedCalc version 23.5.1 (MedCalc Software Ltd., Ostend, Belgium).

## 3 RESULTS

Of the 50 equine cadaver heads (100 TMJs) collected, 9 heads were excluded due to various reasons: wound in the TMJ region penetrating into the joint (n = 2), disruption of the joint capsule (n = 4), fracture of the mandibular condyle (n = 1), and missing of the intra–articular disc sample in the histopathology slides (n = 2). A total of 41 heads (82 TMJs) met the inclusion criteria. These included 23 females and 18 males, all of which were warmblood horses. The median age for the entire horses was 11 years (range 1–25). The age, gender, and sample size for each age category were as follows: (1) 1–4 years (horses n = 17; TMJs n = 34; median age of 2 years (range 1–4); 6 females, 11 males), (2) 5–15 years (horses n = 12; TMJs n = 24; median age of 11.5 years (range 7–15); 8 females, 4 males), and >15 years (horses n = 12; TMJs n = 24; median age of 20 years (range 16–25); 9 females, 3 males).

### 3.1 CT findings in equine TMJs across age–related groups

CT findings were observed in 79% of joints (65/82 TMJs). These findings were partially dichotomously associated with age, with significant differences in the frequency distribution of CT findings between the 1– 4 years and 5–15 years groups (p < 0.0001) as well as between the 1–4 years and >15 years groups (p < 0.0001). However, no significant difference was observed between the 5–15 years and >15 years groups (p = 0.84). A summary of the CT findings is shown in Table 2.

**Table 2.** Distribution of CT findings observed in 82 temporomandibular joints (TMJs) from 41 horses. Horses are divided into three age–related groups. Correlation between each CT finding and the horses’ age is provided in the last column. ‘CT anatomical variations’ are indicated by asterisk.

| CT findings | Age-related groups | | | Correlation<br>($\rho$; $p$) |
| --- | --- | --- | --- | --- |
| | 1–4 years<br>($n = 34$) | 5–15 years<br>($n = 24$) | >15 years<br>($n = 24$) | |
| Bone surface |  |  |  |  |
| Normal | 32 (94%) | 3 (13%) | 0 | $\rho = -0.81$; $p < 0.0001$ |
| Flatt MC* | 0 | 18 (75%) | 24 (100%) | $\rho = 0.85$; $p < 0.0001$ |
| Irr MC | 2 (6%) | 10 (42%) | 8 (33%) | $\rho = 0.35$; $p = 0.001$ |
| Subchondral bone |  |  |  |  |
| Normal | 19 (54%) | 14 (58%) | 15 (63%) | $\rho = 0.10$; $p = 0.37$ |
| ScHD–Scl MC* | 12 (35%) | 0 | 0 | $\rho = -0.50$; $p < 0.0001$ |
| SphHD+Scl MC* | 3 (9%) | 7 (29%) | 7 (29%) | $\rho = 0.37$; $p = 0.0006$ |
| SphHD–Scl MC* | 0 | 3 (13%) | 2 (8%) | $\rho = -0.02$; $p = 0.89$ |
| Bone margins |  |  |  |  |
| Normal | 34 (100%) | 0 | 0 | $\rho = -0.86$; $p < 0.0001$ |
| Medial Ent MC* | 0 | 24 (100%) | 24 (100%) | $\rho = 0.86$; $p < 0.0001$ |
| Lateral Ost MC | 0 | 1 (4%) | 0 | $\rho = 0.07$; $p = 0.52$ |
| Joint space |  |  |  |  |
| Normal | 34 (100%) | 19 (79%) | 21 (88%) | $\rho = -0.17$; $p = 0.12$ |
| Narrow JS | 0 | 5 (21%) | 3 (13%) | $\rho = 0.17$; $p = 0.12$ |
| Joints with findings | 17 (50%) | 24 (100%) | 24 (100%) |  |
| Chi-square test | $p < 0.0001$ | | $p = 0.84$ | |
| | $p < 0.0001$ | | | |

Footnotes: CT = computed tomography; Ent = enthesophytes; Flatt = flattening; Irr = irregularity; JS = joint space; MC = mandibular condyle; n = number of joints with findings; Ost = osteophytes; ScHD = scattered regions of hypodensity; Scl = sclerosis (hyperdensity); SphHD = spherical region of hypodensity; % = percentage of joints with findings from each age-related group; \* = 'CT anatomical variations'. The differences in the frequency distribution of CT findings between age-related groups were considered significant at $p < 0.05$ . The Spearman correlation coefficient ( $\rho$ ) was considered significant at $p < 0.05$ and then highlighted with bold font.

In 1–4 years group, CT findings were observed in 50% of joints (17/34 TMJs), while in the remaining 50% (17/34 TMJs), the mandibular condyles were globose and homogeneous (Figure 1A). In this age–related group, irregularity of the mandibular condyle was observed only in 6% of joints (Figure 1B). Scattered regions of hypodensity (Figure 1C) and spherical regions of hypodensity with surrounding sclerosis (Figure 1D) were observed in the subchondral bone of the mandibular condyle in 12% and 9% of joints, respectively. Spherical regions of hypodensity in two joints were close to the articular surface of the bone, and in one joint bone cyst was open to the articular surface of the bone. In this age–related group, 15% of the entire CT findings were limited to the right joint, 6% to the left joint, and 29% were bilateral.

**Figure 1.**
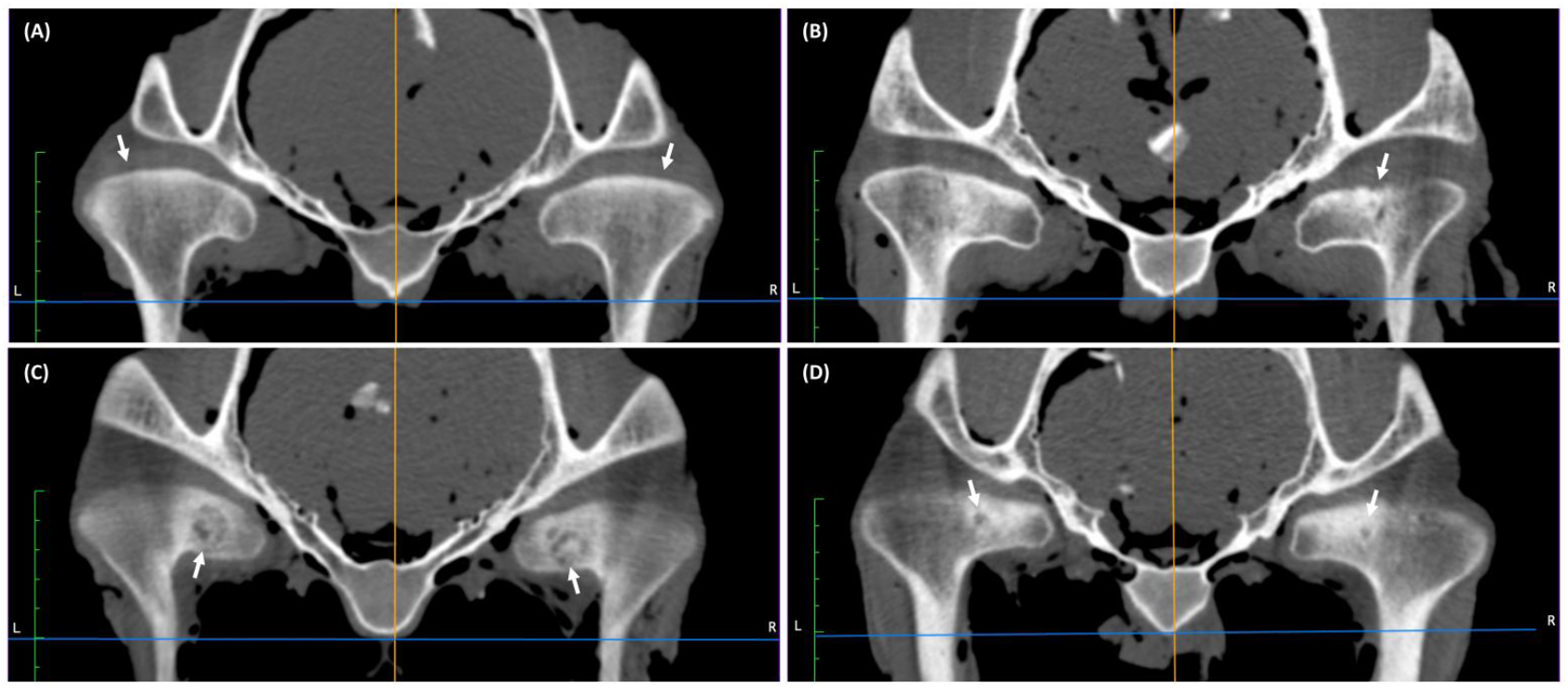
Examples of transverse computed tomography (CT) images of temporomandibular joints (TMJs) in cadaver heads of horses aged 1–4 years. A, Globose shape (white arrows) and homogenous mandibular condyles in joints without CT findings. B, Irregularity of the bone surface of the right mandibular condyles (white arrow). C, Scattered regions of hypodensity in the right and left mandibular condyles (white arrows). D, Small spherical region of hypodensity (bone cysts) with surrounding sclerosis in the right and left mandibular condyles (white arrows).

In 5–15 years group, CT findings were observed in 100% of joints (24/24 TMJs). In this age–related group, flattening (Figure 2A-D) and irregularity of the mandibular condyle (Figure 2B,D) were observed in 75% and 42% of joints, respectively. Spherical regions of hypodensity (Figure 2C) in the mandibular condyle were found in 29% of joints with surrounding sclerosis, and in 13% of joints without. Spherical regions of hypodensity in seven joints were close to the articular surface of the bone, and in three joints were open to the articular surface of the bone. Medial enthesophytes were observed in 100% of joints (Figure 2A-D), whereas lateral osteophytes only in 4% of joints (Figure 2D). Joint space narrowing was detected in 21% of joints (Figure 2D). CT findings were observed bilaterally in all joints.

**Figure 2.**
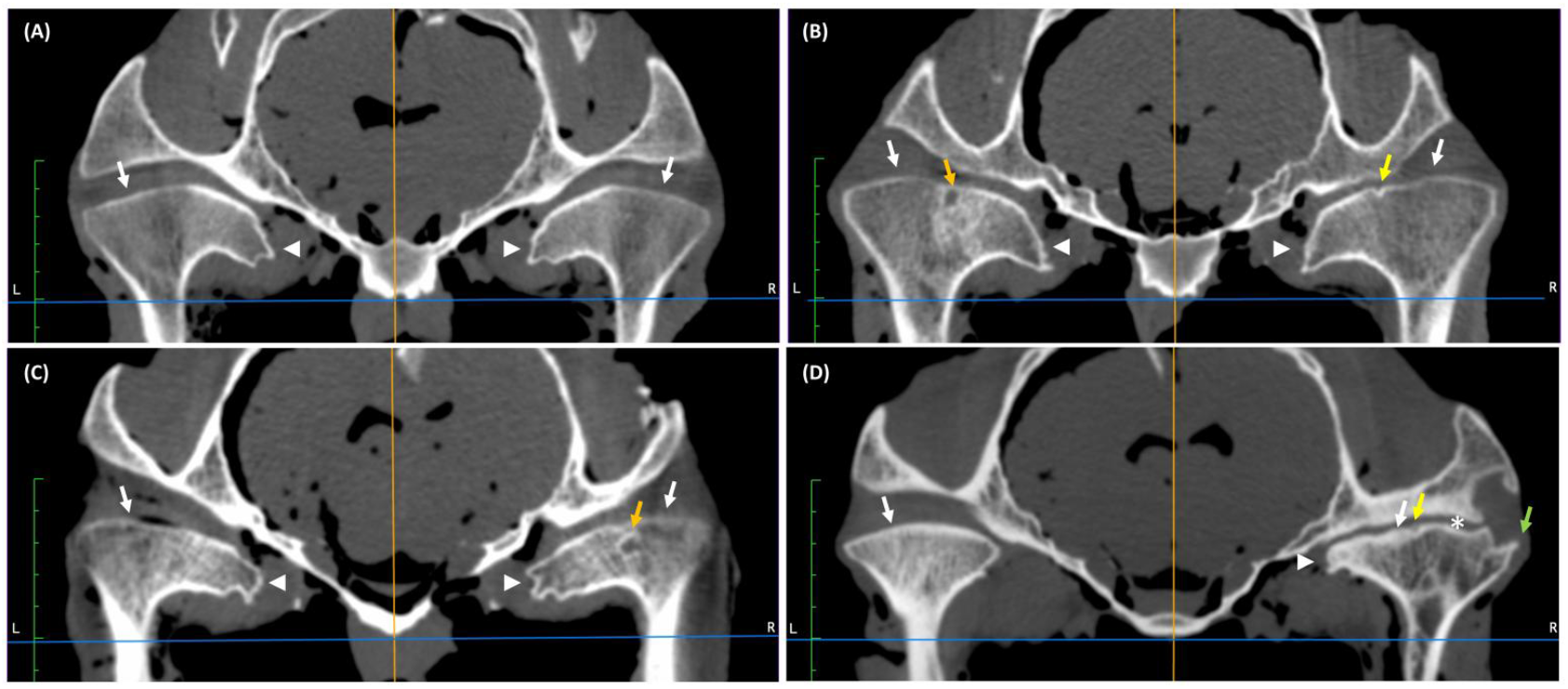
Examples of transverse computed tomography (CT) images of temporomandibular joints (TMJs) in cadaver heads of horses aged 5–15 years. A, Flattening (white arrows) and medial enthesophytes (white arrow heads) of both mandibular condyles. B, Irregularity of the right mandibular condyle (yellow arrow) and spherical region of hypodensity (bone cysts) with surrounding sclerosis in the left mandibular condyle (orange arrow). Flattening (white arrows) and medial enthesophytes of both mandibular condyles (white arrow heads). C, Spherical region of hypodensity (bone cysts) with surrounding sclerosis in the caudal aspect of right mandibular condyle (orange arrow). Flattening (white arrows) and medial enthesophytes (white arrow heads) of both mandibular condyles. D, Irregularity (yellow arrow), medial enthesophytes (white arrow head), and lateral osteophytes (green arrow) of the right mandibular condyle. Narrowing of the right joint space (asterisk). Flattening (white arrows) of both mandibular condyles.

In >15 years group, CT findings were observed in 100% of joints (24/24 TMJs). In this age–related group, flattening (Figure 3A-D) and irregularity (Figure 3B) of the mandibular condyle were observed in 100% and 33% of joints, respectively. Spherical regions of hypodensity in the mandibular condyle were found in 29% of joints with surrounding sclerosis (Figure 3D) and in 8% of joints without (Figure 3C). Spherical regions of hypodensity in five joints were close to the articular surface of the bone, whereas in four joints were open to the articular surface of the bone. Medial enthesophytes were noted in 100% of joints (Figure 3A-D), and joint space narrowing was detected in 21% of joints (Figure 3D). CT findings were observed bilaterally in all joints.

**Figure 3.**
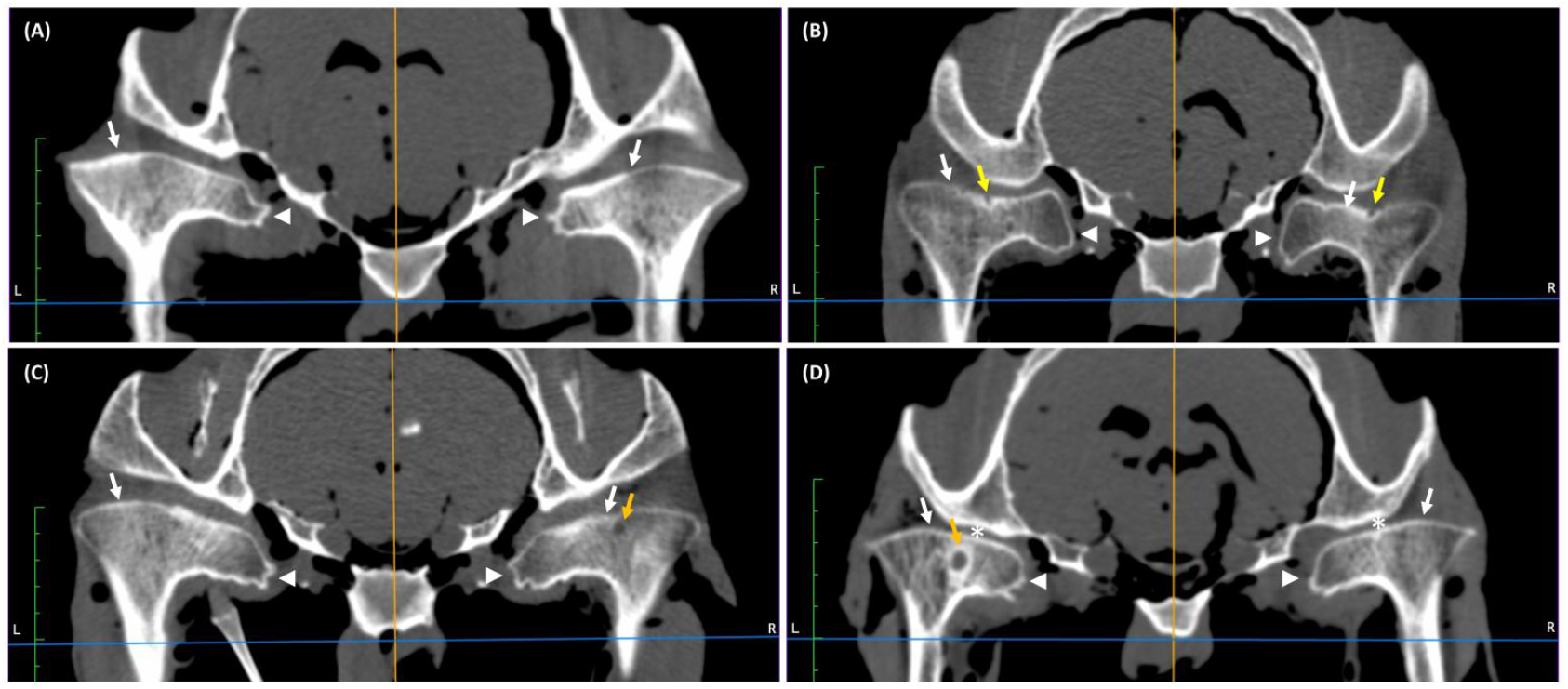
Examples of transverse computed tomography (CT) images of temporomandibular joints (TMJs) in cadaver heads of horses aged >15 years. A, Flattening (white arrows) and medial enthesophytes (white arrow heads) of both mandibular condyles. B, Irregularity of both mandibular condyles (yellow arrows). Flattening (white arrows) and medial enthesophytes (white arrow heads) of both mandibular condyles. C, Spherical region of hypodensity (bone cysts) without surrounding sclerosis in the right mandibular condyle (orange arrow). Flattening (white arrows) and medial enthesophytes (white arrow heads) of both mandibular condyles. D, Spherical region of hypodensity (bone cysts) with surrounding sclerosis in the left mandibular condyle (orange arrow). Flattening (white arrows) and medial enthesophytes (white arrow heads) of both mandibular condyles. Narrowing of both joint spaces (asterisks).

A strong positive correlations were found between age and flattening of the mandibular condyle (ρ = 0.85; p < 0.0001), as well as between age and medial enthesophytes on its margin (ρ = 0.86; p < 0.0001). A moderate negative correlation was found between age and scattered hypodensity in the mandibular condyle (ρ = -0.50; p < 0.0001). A weak positive correlations were noted between age and irregularity of the mandibular condyle (ρ = 0.35; p = 0.001), as well as age and spherical regions of hypodensity with surrounding sclerosis in the mandibular condyle (ρ = 0.37; p = 0.0006). In contrast, strong negative correlations were found between age and normal CT appearance of the bone surface of the mandibular condyle (ρ = -0.81; p < 0.0001), as well as age and normal CT appearance of the bone margins (ρ = -0.86; p < 0.0001).

### 3.2 Histopathological findings in equine TMJs across age–related groups

Histopathological findings were observed in 50% of joints (41/82 TMJs) and showed a clear dichotomous age–related association. The frequency distribution of histopathological findings differed between the 1–4 years and 5–15 years groups (p < 0.0001), the 1–4 years and >15 years groups (p < 0.0001), as well as the 5–15 years and >15 years groups (p < 0.0001). A summary of histopathological findings is shown in Table 3.

**Table 3.**
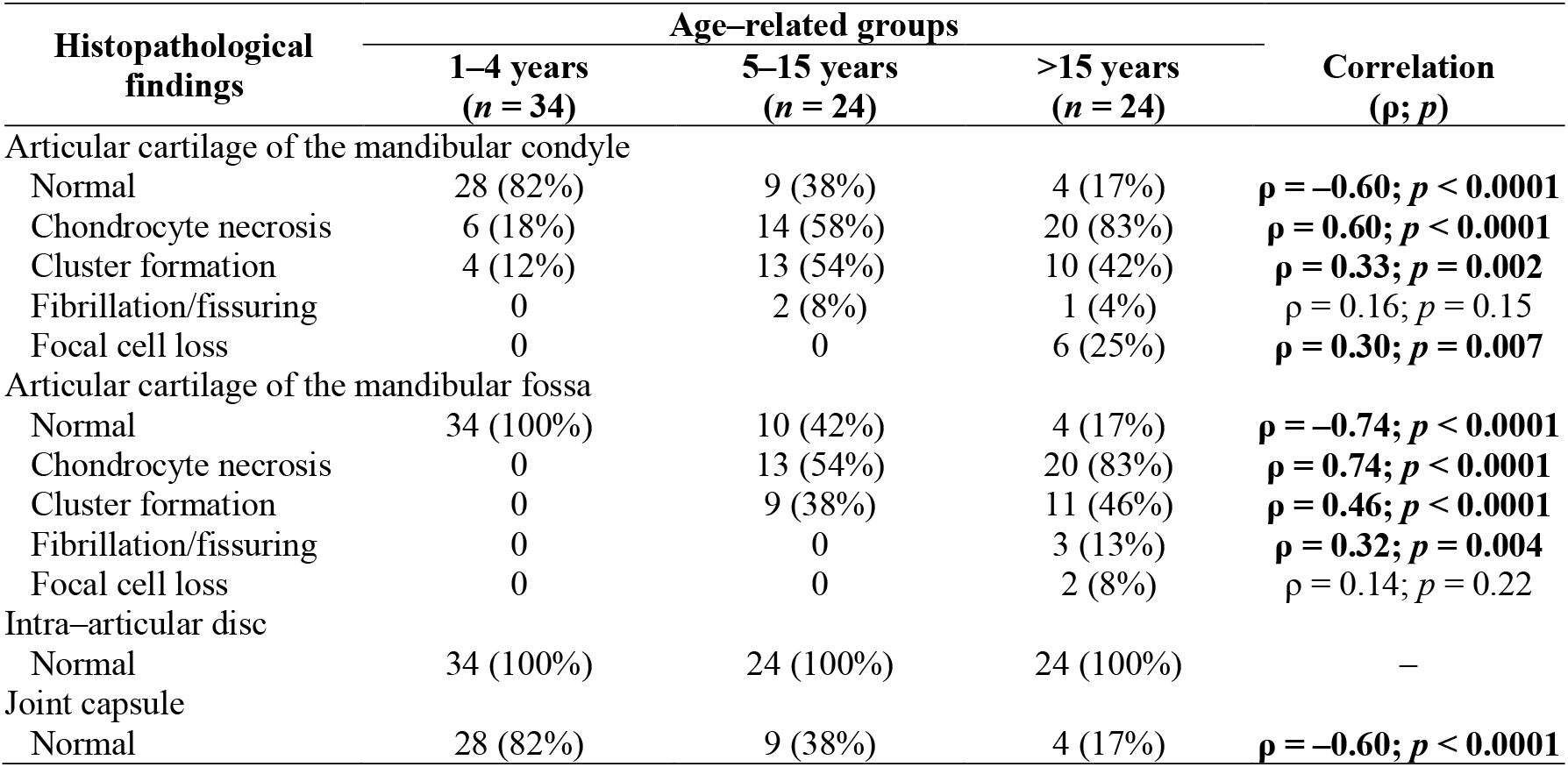
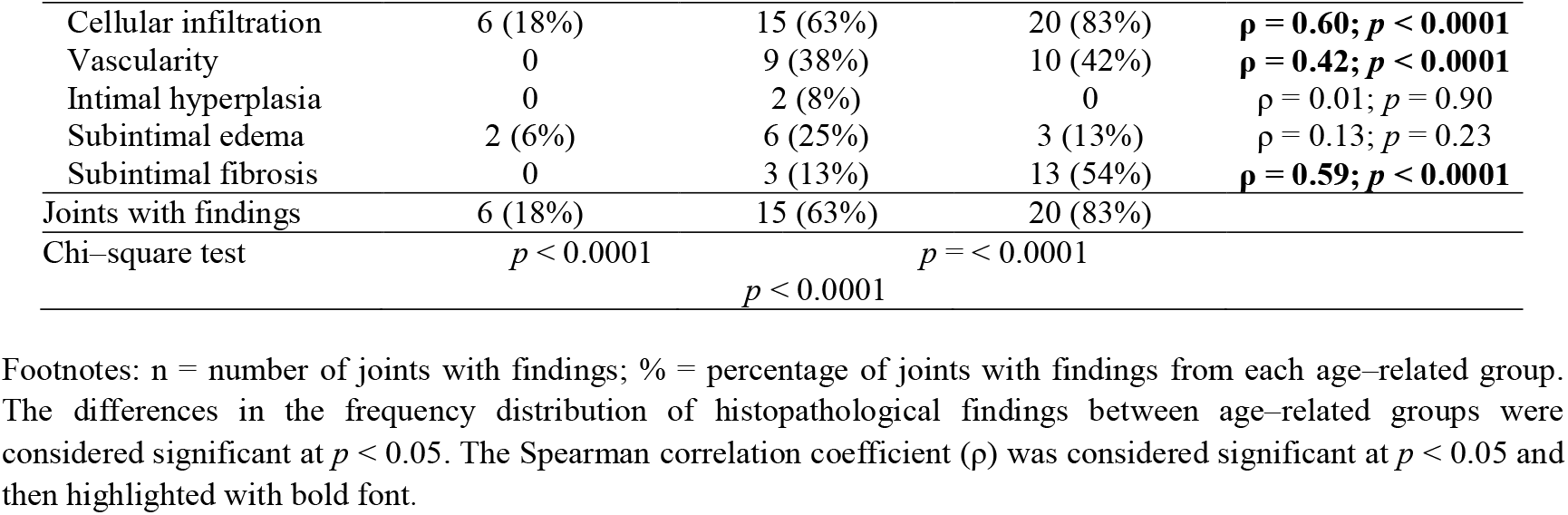
Distribution of histopathological findings observed in 82 temporomandibular joints (TMJs) from 41 horses. Horses are divided into three age–related groups. Correlation between each histopathological finding and the horses’ age is provided in the last column.

In 1–4 years group, histopathological findings were observed in 18% of joints (6/34 TMJs), whereas no histopathological abnormalities were identified in the remaining 82% (28/34 TMJs) in either the mandibular condyle (Figure 4A), the mandibular fossa (Figure 4B), or the joint capsule (Figure 4C). In this age–related group, chondrocyte necrosis and cluster formation in the mandibular condyle articular cartilage were found in 18% and 12% of joints, respectively. Cellular infiltration and subintimal edema in joint capsule were presented in 18% and 6% of joints, respectively (Figure 4D). Of those horses, 12% of findings were limited to the right joint, whereas 6% were bilateral.

**Figure 4.**
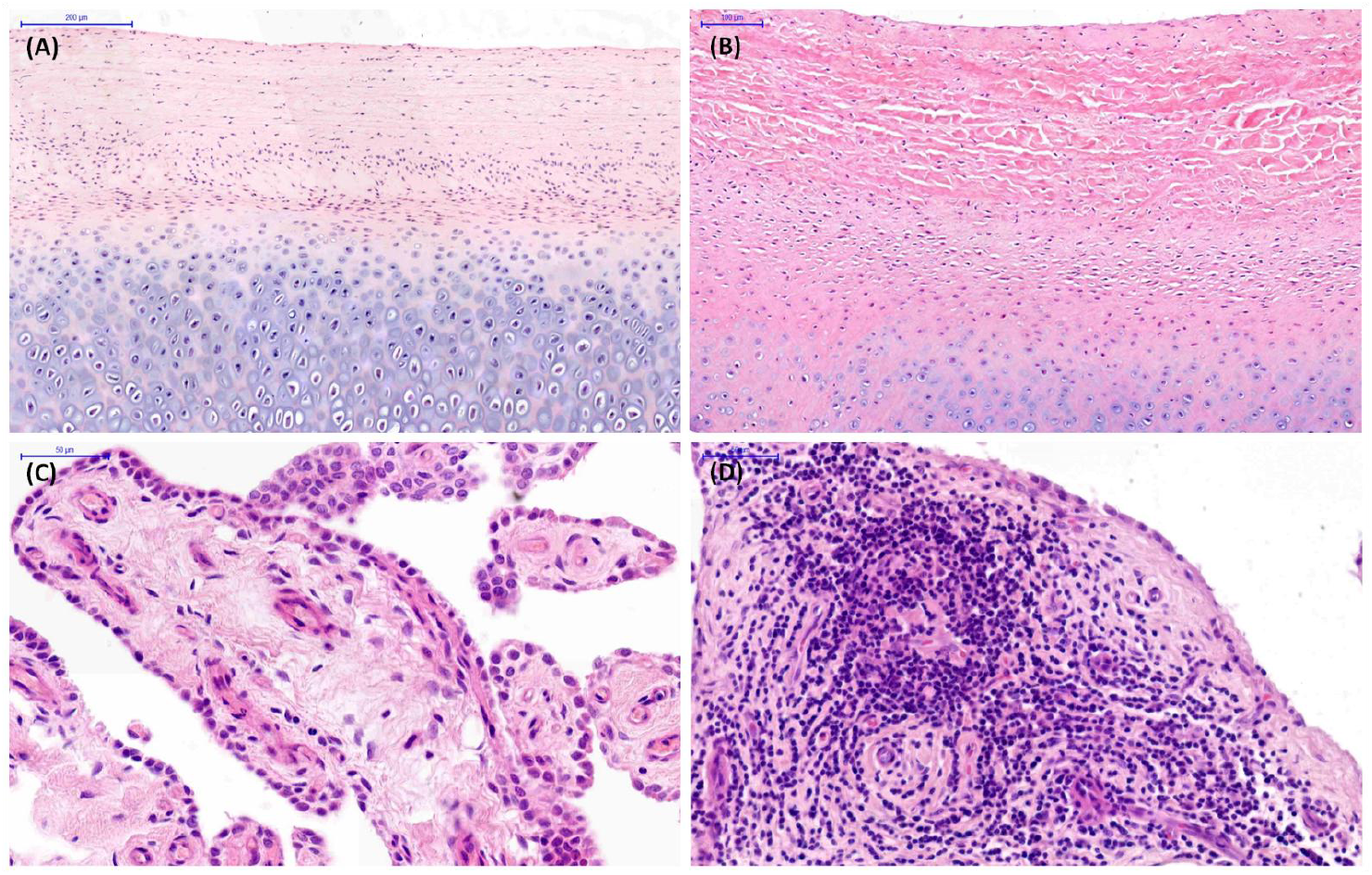
Examples of histopathological images of temporomandibular joint (TMJ) tissue sections collected from cadaver heads of horses aged 1–4 years. A, Normal articular cartilage in the mandibular condyle. B, Normal articular cartilage in the mandibular fossa. C, Normal joint capsule. D, Cellular infiltration in the joint capsule.

In 5–15 years group, histopathological findings were observed in 63% of joints (15/24 TMJs). In this age–related group, chondrocyte necrosis, cluster formation, and fibrillation/fissuring of the mandibular condyle articular cartilage were found in 58%, 54%, and 8% of joints, respectively (Figure 5A). Chondrocyte necrosis and cluster formation in the mandibular fossa articular cartilage were observed in 54% and 38% of joints, respectively (Figure 5B). Vascularity, subintimal edema, subintimal fibrosis, and intimal hyperplasia were noted in 38%, 25%, 13%, and 8% of joints, respectively (Figure 5C); whereas cellular infiltration in joint capsule was presented in 63% of joints (Figure 5D). Histopathological findings were bilateral in 58% of joints, and in 4% were limited to the left joint.

**Figure 5.**
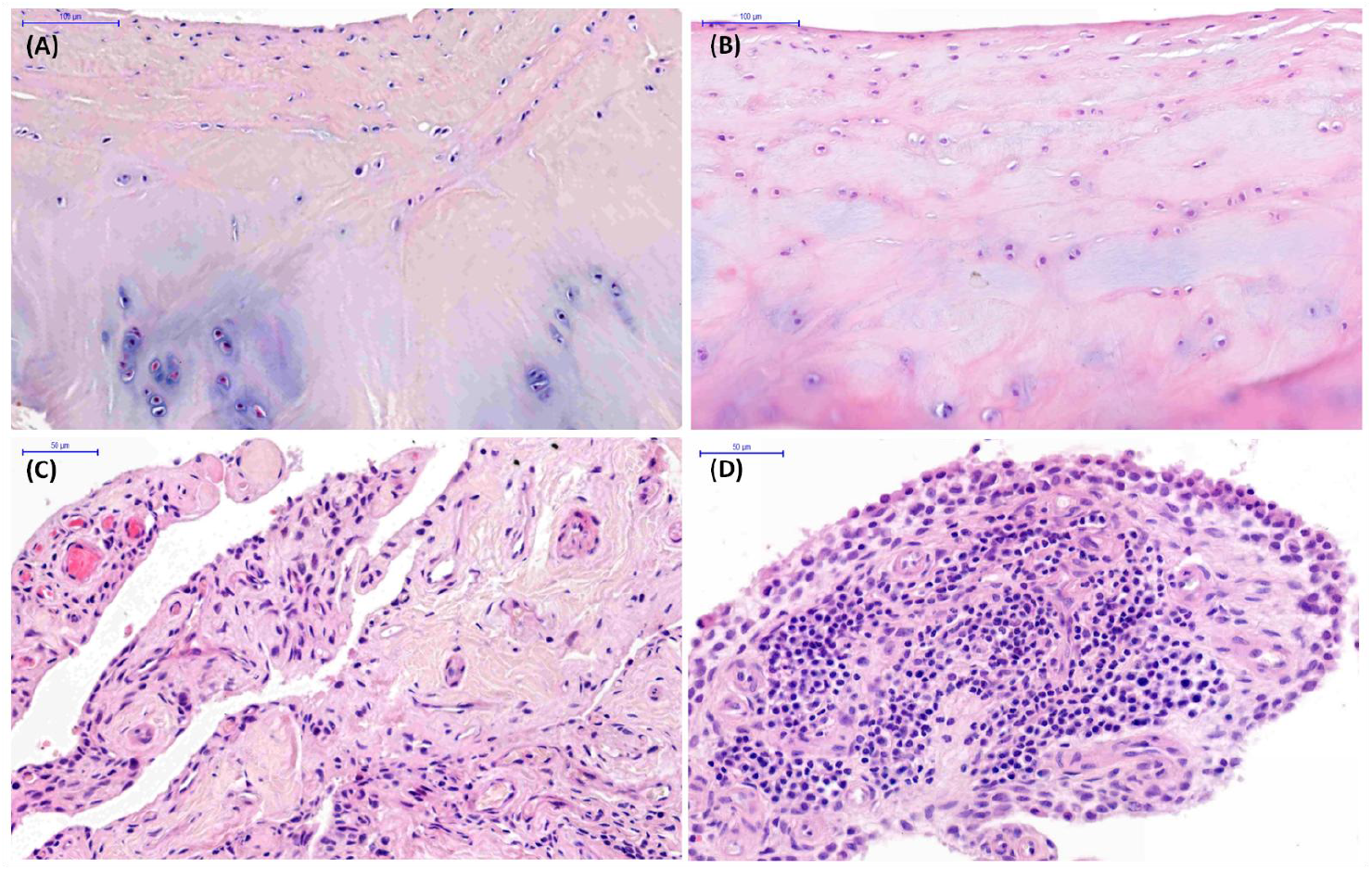
Examples of histopathological images of temporomandibular joint (TMJ) tissue sections collected from cadaver heads of horses aged 5–15 years. A, Chondrocyte necrosis and cluster formation in the mandibular condyle articular cartilage. B, Chondrocyte necrosis and cluster formation in the mandibular fossa articular cartilage. C, Vascularity and subintimal edema in the joint capsule. D, Cellular infiltration in the joint capsule.

In >15 years group, histopathological findings were observed in 83% of joints (20/24 TMJs). In this age–related group, chondrocyte necrosis, cluster formation, focal cell loss, and fibrillation/fissuring in the mandibular condyle articular cartilage were found in 83%, 42%, 25%, and 4% of joints, respectively (Figure 6A). Chondrocyte necrosis, cluster formation, focal cell loss, and fibrillation/fissuring in the mandibular fossa articular cartilage were observed in 83%, 48%, 8%, and 13% of joints, respectively (Figure 6A). Subintimal fibrosis, vascularity, and subintimal edema were noted in 54%, 42%, and 13% of joints, respectively (Figure 6C); whereas cellular infiltration in joint capsule was presented in 83% of joints (Figure 6D). Bilateral histopathological findings were presented in 75% of joints, and 8% were limited to the right joint.

**Figure 6.**
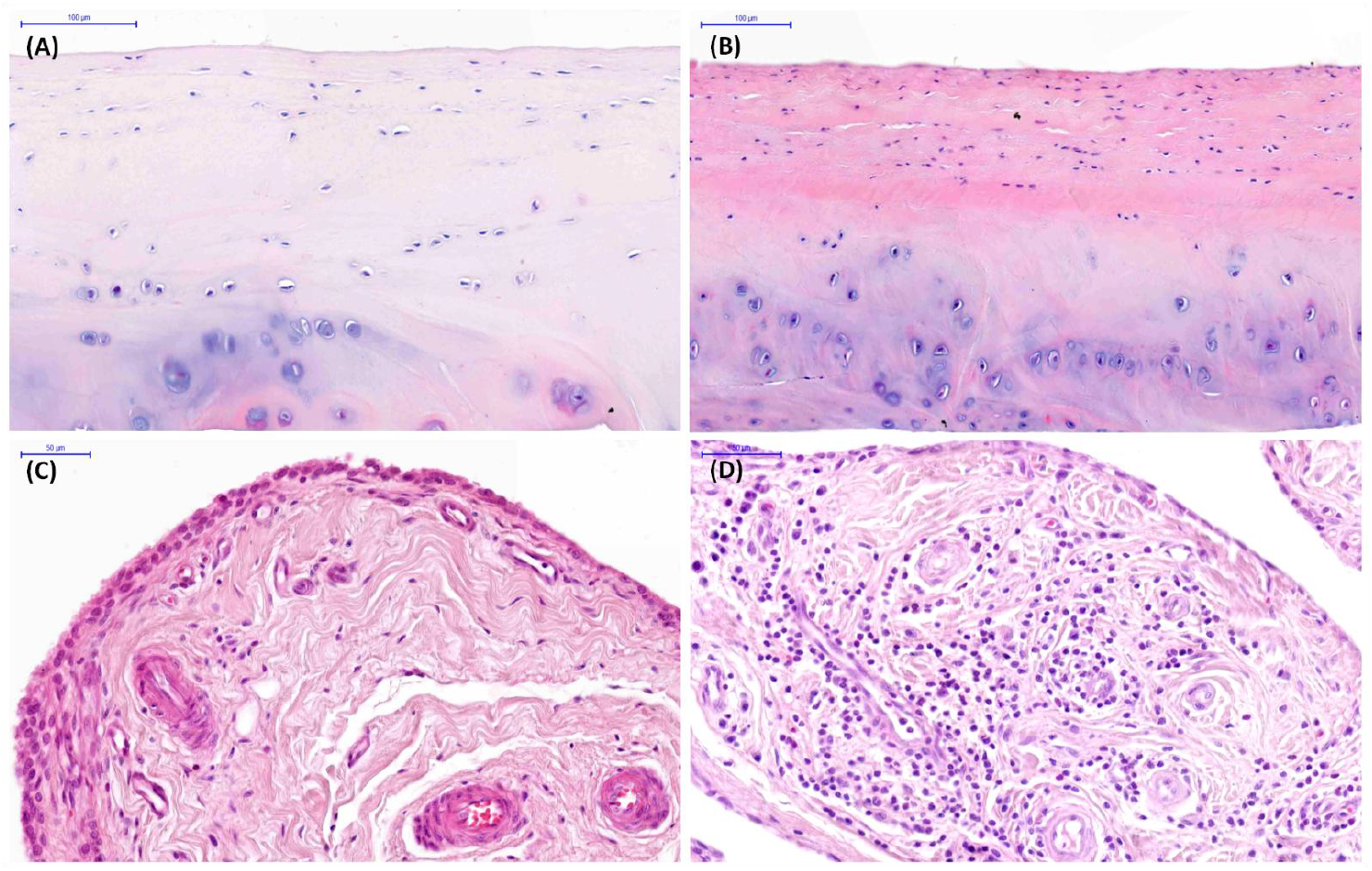
Examples of histopathological images of temporomandibular joint (TMJ) tissue sections collected from cadaver heads of horses aged >15 years. A, Chondrocyte necrosis, cluster formation, and focal cell loss in the mandibular condyle articular cartilage. B, Chondrocyte necrosis in the mandibular fossa articular cartilage. C, Subintimal fibrosis in the joint capsule. D, Cellular infiltration and vascularity in the joint capsule.

A strong positive correlation was found between age and chondrocyte necrosis in the mandibular fossa bone articular cartilage (ρ = 0.85; p < 0.0001). In the articular cartilage, moderate positive correlations were observed between age and chondrocyte necrosis in the mandibular condyle (ρ = 0.60; p < 0.0001), as well as between age and cluster formation in the mandibular fossa (ρ = 0.46; p < 0.0001). In the joint capsule, moderate positive correlations were noted between age and cellular infiltration (ρ = 0.60; p < 0.0001), subintimal fibrosis (ρ = 0.59; p < 0.0001), and vascularity (ρ = 0.42; p < 0.0001). Weak positive correlations was found in the articular cartilage between age and cluster formation in the mandibular condyle (ρ = 0.33; p = 0.002), focal cell loss in the mandibular condyle (ρ = 0.30; p = 0.007), and fibrillation/fissuring in the mandibular fossa (ρ = 0.32; p = 0.004). In contrast, moderate to strong negative correlations were observed between age and normal histopathological appearance in the mandibular condyle articular cartilage (ρ = -0.60; p < 0.0001), mandibular fossa articular cartilage (ρ = -0.74; p < 0.0001), as well as joint capsule (ρ = -0.60; p < 0.0001)

### 3.3 Relationship Between CT and histopathological findings in equine TMJs

The correlations between CT and histopathological findings are summarized in Table 4. A strong positive correlation was observed between flattening of the mandibular condyle and chondrocyte necrosis in the mandibular fossa articular cartilage (ρ = 0.75; p < 0.0001). Moderate positive correlations were identified between flattening of the mandibular condyle and chondrocyte necrosis in the mandibular condyle articular cartilage (ρ = 0.56; p < 0.0001), cluster formation in the mandibular fossa articular cartilage (ρ = 0.44; p < 0.0001), and two alterations in the joint capsule, including cellular infiltration (ρ = 0.54; p < 0.0001) and vascularity (ρ = 0.48; p < 0.0001). Moderate positive correlations were also found between irregularity of the mandibular condyle and chondrocyte necrosis in both the mandibular condyle (ρ = 0.53; p < 0.0001) and mandibular fossa (ρ = 0.46; p < 0.0001) articular cartilage; cluster formation also in both the mandibular condyle (ρ = 0.45; p < 0.0001) and mandibular fossa (ρ = 0.40; p = 0.0002) articular cartilage; as well as cellular infiltration in the joint capsule (ρ = 0.57; p < 0.0001). Moderate positive correlations were observed between spherical regions of hypodensity with surrounding sclerosis in the mandibular condyle and chondrocyte necrosis in both the mandibular condyle (ρ = 0.52; p < 0.0001) and mandibular fossa (ρ = 0.44; p < 0.0001) articular cartilage; cluster formation in the mandibular condyle articular cartilage (ρ = 0.47; p < 0.0001); as well as cellular infiltration in the joint capsule (ρ = 0.51; p < 0.0001). Furthermore, moderate positive correlations were found between medial enthesophytes and chondrocyte necrosis in both the mandibular condyle (ρ = 0.52; p < 0.0001) and mandibular fossa (ρ = 0.69; p < 0.0001) articular cartilage, cluster formation in the mandibular fossa articular cartilage (ρ = 0.48; p < 0.0001), and three alterations in the joint capsule, including cellular infiltration (ρ = 0.54; p < 0.0001), vascularity (ρ = 0.46; p < 0.0001), and subintimal fibrosis (ρ = 0.41; p = 0.0001).

**Table 4.** The Spearman correlation coefficient (ρ) between CT and histopathological findings observed in 82 temporomandibular joints (TMJs) from 41 horses. ‘CT anatomical variations’ are indicated by asterisk.

| CT findings /<br>histopathological<br>findings | Bone surface |  | Subchondral bone |  |  | Bone margins |  | Joint space |
| --- | --- | --- | --- | --- | --- | --- | --- | --- |
|  | Flatt MC* | Irr MC | ScHD–Scl<br>MC* | SphHD+Scl<br>MC* | SphHD–Scl<br>MC* | Medial Ent<br>MC* | Lateral Ost<br>MC | Narrow JS |
| Articular cartilage of the mandibular condyle |  |  |  |  |  |  |  |  |
| Chondrocyte necrosis | $\rho = 0.56$;<br>$p < 0.0001$ | $\rho = 0.53$;<br>$p < 0.0001$ | $\rho = -0.34$;<br>$p = 0.002$ | $\rho = 0.52$;<br>$p < 0.0001$ | $\rho = -0.04$;<br>$p = 0.69$ | $\rho = 0.52$;<br>$p < 0.0001$ | $\rho = 0.11$;<br>$p = 0.31$ | $\rho = 0.01$;<br>$p = 0.94$ |
| Cluster formation | $\rho = 0.32$;<br>$p = 0.003$ | $\rho = 0.45$;<br>$p < 0.0001$ | $\rho = -0.22$;<br>$p = 0.05$ | $\rho = 0.47$;<br>$p < 0.0001$ | $\rho = -0.07$;<br>$p = 0.53$ | $\rho = 0.38$;<br>$p = 0.0004$ | $\rho = 0.16$;<br>$p = 0.15$ | $\rho = 0.12$;<br>$p = 0.29$ |
| Fibrillation/fissuring | $\rho = 0.06$;<br>$p = 0.59$ | $\rho = 0.04$;<br>$p = 0.72$ | $\rho = -0.08$;<br>$p = 0.47$ | $\rho = 0.22$;<br>$p = 0.046$ | $\rho = -0.05$;<br>$p = 0.66$ | $\rho = 0.16$;<br>$p = 0.14$ | $\rho = -0.02$;<br>$p = 0.85$ | $\rho = -0.06$;<br>$p = 0.57$ |
| Focal cell loss | $\rho = 0.27$;<br>$p = 0.01$ | $\rho = 0.06$;<br>$p = 0.60$ | $\rho = -0.12$;<br>$p = 0.30$ | $\rho = 0.20$;<br>$p = 0.07$ | $\rho = -0.07$;<br>$p = 0.52$ | $\rho = 0.24$;<br>$p = 0.03$ | $\rho = -0.03$;<br>$p = 0.78$ | $\rho = 0.07$;<br>$p = 0.56$ |
| Articular cartilage of the mandibular fossa |  |  |  |  |  |  |  |  |
| Chondrocyte necrosis | $\rho = 0.75$;<br>$p < 0.0001$ | $\rho = 0.46$;<br>$p < 0.0001$ | $\rho = -0.34$;<br>$p = 0.002$ | $\rho = 0.44$;<br>$p < 0.0001$ | $\rho = -0.001$;<br>$p = 0.99$ | $\rho = 0.69$;<br>$p < 0.0001$ | $\rho = 0.14$;<br>$p = 0.23$ | $\rho = 0.07$;<br>$p = 0.56$ |
| Cluster formation | $\rho = 0.44$;<br>$p < 0.0001$ | $\rho = 0.40$;<br>$p = 0.0002$ | $\rho = -0.24$;<br>$p = 0.03$ | $\rho = 0.27$;<br>$p = 0.01$ | $\rho = -0.03$;<br>$p = 0.82$ | $\rho = 0.48$;<br>$p < 0.0001$ | $\rho = 0.20$;<br>$p = 0.08$ | $\rho = 0.10$;<br>$p = 0.37$ |
| Fibrillation/fissuring | $\rho = 0.19$;<br>$p = 0.09$ | $\rho = 0.19$;<br>$p = 0.08$ | $\rho = -0.08$;<br>$p = 0.47$ | $\rho = 0.22$;<br>$p = 0.046$ | $\rho = -0.05$;<br>$p = 0.66$ | $\rho = 0.16$;<br>$p = 0.14$ | $\rho = -0.02$;<br>$p = 0.85$ | $\rho = -0.06$;<br>$p = 0.57$ |
| Focal cell loss | $\rho = 0.15$;<br>$p = 0.17$ | $\rho = 0.09$;<br>$p = 0.40$ | $\rho = -0.07$;<br>$p = 0.56$ | $\rho = 0.11$;<br>$p = 0.31$ | $\rho = -0.05$;<br>$p = 0.72$ | $\rho = 0.16$;<br>$p = 0.23$ | $\rho = -0.02$;<br>$p = 0.88$ | $\rho = -0.06$;<br>$p = 0.64$ |
| Joint capsule |  |  |  |  |  |  |  |  |
| Cellular infiltration | $\rho = 0.54$;<br>$p < 0.0001$ | $\rho = 0.57$;<br>$p < 0.0001$ | $\rho = -0.35$;<br>$p = 0.002$ | $\rho = 0.51$;<br>$p < 0.0001$ | $\rho = -0.05$;<br>$p = 0.65$ | $\rho = 0.54$;<br>$p < 0.0001$ | $\rho = 0.11$;<br>$p = 0.32$ | $\rho = 0.08$;<br>$p = 0.46$ |
| Vascularity | $\rho = 0.48$;<br>$p < 0.0001$ | $\rho = 0.23$;<br>$p = 0.04$ | $\rho = -0.23$;<br>$p = 0.04$ | $\rho = 0.36$;<br>$p = 0.001$ | $\rho = -0.14$;<br>$p = 0.21$ | $\rho = 0.46$;<br>$p < 0.0001$ | $\rho = 0.20$;<br>$p = 0.07$ | $\rho = 0.01$;<br>$p = 0.90$ |
| Intimal hyperplasia | $\rho = -0.004$;<br>$p = 0.97$ | $\rho = 0.28$;<br>$p = 0.01$ | $\rho = -0.07$;<br>$p = 0.56$ | $\rho = 0.11$;<br>$p = 0.31$ | $\rho = -0.04$;<br>$p = 0.72$ | $\rho = 0.13$;<br>$p = 0.23$ | $\rho = -0.02$;<br>$p = 0.88$ | $\rho = -0.05$;<br>$p = 0.64$ |
| Subintimal edema | $\rho = 0.24$;<br>$p = 0.03$ | $\rho = 0.28$;<br>$p = 0.01$ | $\rho = -0.16$;<br>$p = 0.14$ | $\rho = 0.24$;<br>$p = 0.03$ | $\rho = -0.05$;<br>$p = 0.66$ | $\rho = 0.19$;<br>$p = 0.09$ | $\rho = 0.28$;<br>$p = 0.01$ | $\rho = 0.23$;<br>$p = 0.04$ |
| Subintimal fibrosis | $\rho = 0.36$;<br>$p = 0.001$ | $\rho = 0.37$;<br>$p = 0.001$ | $\rho = -0.20$;<br>$p = 0.07$ | $\rho = 0.28$;<br>$p = 0.01$ | $\rho = 0.003$;<br>$p = 0.98$ | $\rho = 0.41$;<br>$p = 0.0001$ | $\rho = -0.05$;<br>$p = 0.63$ | $\rho = -0.06$;<br>$p = 0.60$ |

Footnotes: CT = computed tomography; Ent = enthesophytes; Flatt = flattening; Irr = irregularity; JS = joint space; MC = mandibular condyle; Ost = osteophytes; ScHD = scattered regions of hypodensity; Scl = sclerosis (hyperdensity); SphHD = spherical region of hypodensity; + = with; – = without; \* = 'CT anatomical variations'. The correlations were considered significant at $p < 0.05$ and then highlighted with bold font.

Weak positive correlations were identified between flattening of the mandibular condyle and two alterations in the mandibular condyle articular cartilage, including cluster formation (ρ = 0.32; p = 0.003) and focal cell loss (ρ = 0.27; p = 0.01); as well as two alterations in the joint capsule, including subintimal edema (ρ = 0.24; p = 0.03) and subintimal fibrosis (ρ = 0.36; p = 0.001). Weak positive correlations were also found between irregularity of the mandibular condyle and four alterations in the joint capsule, including vascularity (ρ = 0.23; p = 0.04), intimal hyperplasia (ρ = 0.28; p = 0.01), subintimal edema (ρ = 0.28; p = 0.01), and subintimal fibrosis (ρ = 0.37; p = 0.001). Weak positive correlations were observed between spherical regions of hypodensity with surrounding sclerosis in the mandibular condyle and two alteration in the mandibular condyle articular cartilage, including fibrillation/fissuring (ρ = 0.22; p = 0.046) and focal cell loss (ρ = 0.20; p = 0.07).

These spherical regions of hypodensity in the mandibular condyle were also weakly positively correlated with cluster formation (ρ = 0.27; p = 0.01) and fibrillation/fissuring (ρ = 0.22; p = 0.046) in the mandibular fossa articular cartilage; as well as vascularity (ρ = 0.36; p = 0.001), subintimal edema (ρ = 0.24; p = 0.03), and subintimal fibrosis (ρ = 0.28; p = 0.01) in the joint capsule. Moreover, weak positive correlations were found between medial enthesophytes and two alterations in the mandibular condyle articular cartilage, including cluster formation (ρ = 0.38; p = 0.0004) and focal cell loss (ρ = 0.24; p = 0.03), lateral osteophytes and subintimal edema in the joint capsule (ρ = 0.28; p = 0.01), as well as joint space narrowing and subintimal edema in the joint capsule (ρ = 0.23; p = 0.04). Weak negative correlations were identified between scattered regions of hypodensity in the mandibular condyle and chondrocyte necrosis in both the mandibular condyle (ρ = -0.34; p = 0.002) and mandibular fossa (ρ = -0.34; p = 0.002) articular cartilage, cluster formation in the mandibular fossa articular cartilage (ρ = -0.24; p = 0.03), as well as two alterations in the joint capsule, including cellular infiltration (ρ = -0.35; p = 0.002) and vascularity (ρ = - 0.23; p = 0.04).

Distribution of the CT findings among TMJ OA–related groups is shown in Table 5. CT findings were noted in 100% of joints (41/41 TMJs) with histopathologically confirmed OA and in 59% of joints (24/41 TMJs) showing no histopathological findings of (OA–free). The frequency distribution of CT findings differed between the TMJ OA and OA–free TMJ groups (p < 0.0001), with no clear dichotomy regarding histopathological confirmation of OA.

**Table 5.** Distribution of CT findings observed in 82 temporomandibular joints (TMJs) from 41 horses among two TMJ osteoarthritis (OA)–related groups, including group with histopathologically confirmed TMJ OA group and OA– free TMJ group. ‘CT anatomical variations’ are indicated by asterisk.

| CT findings | TMJ OA-related groups |  |
| --- | --- | --- |
|  | TMJ OA group<br>(n = 41) | OA-free TMJ group<br>(n = 41) |
| Bone surface |  |  |
| Normal | 4 (10%) | 31 (76%) |
| Flatt MC* | 32 (78%) | 10 (24%) |
| Irr MC | 20 (49%) | 0 |
| Subchondral bone |  |  |
| Normal | 21 (51%) | 27 (66%) |
| ScHD-Scl MC* | 1 (2%) | 11 (27%) |
| SphHD+Scl MC* | 17 (41%) | 0 |
| SphHD-Scl MC* | 2 (5%) | 3 (7%) |
| Bone margins |  |  |
| Normal | 6 (15%) | 28 (68%) |
| Medial Ent MC* | 35 (85%) | 13 (32%) |
| Lateral Ost MC | 1 (2%) | 0 |
| Joint space |  |  |
| Normal | 36 (88%) | 38 (93%) |
| Narrow JS | 5 (12%) | 3 (7%) |
| Joints with findings | 41 (100%) | 24 (59%) |
| Chi-square test | $p < 0.0001$ | |

Footnotes: CT = computed tomography; Ent = enthesophytes; Flatt = flattening; Irr = irregularity; JS = joint space; MC = mandibular condyle; n = number of joints with findings; Ost = osteophytes; ScHD = scattered regions of hypodensity; Scl = sclerosis (hyperdensity); SphHD = spherical region of hypodensity; % = percentage of joints with findings; \* = 'CT anatomical variations'. The differences in the frequency distribution of CT findings between TMJ OA-related groups were considered significant at $p < 0.05$ .

In the TMJ OA group, flattening and irregularity of the mandibular condyle were observed in 78% and 49% of joints, respectively. In the subchondral bone of mandibular condyle, scattered regions of hypodensity were noted in 2% of joints, whereas spherical regions of hypodensity were observed in 46% of joints, including bone cysts with surrounding sclerosis in 41% of joints and bone cysts without surrounding sclerosis in 5% of joints. Medial enthesophytes were identified in 85% of joints, while lateral osteophytes were presented in 2% of joints. Joint space narrowing was detected in 12% of joints. Notably, in TMJ OA group, the absence of CT findings was observed in 10% of joints considering findings in bone surface, 51% of joints considering findings in subchondral bone, 15% of joints considering findings in bony margins, and 88% in joint space.

In the OA–free TMJ group, the absence of CT findings was observed in a proportion of joints: 76% for findings in bone surface, 66% for findings in subchondral bone, 68% for findings in bony margins, and 93% for findings in joint space. Notably, in the OA–free TMJ group, the flattening of the mandibular condyle was still observed in 24% of joints. In the subchondral bone of mandibular condyle, scattered regions of hypodensity were noted in 27% of joints, whereas spherical regions of hypodensity without surrounding sclerosis were observed 7% of joints. Medial enthesophytes were identified in 32% of joints, and joint space narrowing was detected in 7% of joints.

### 3.4 Accuracy of CT in diagnosing TMJ OA

The accuracy metrics are summarized shown in Table 6. In subgroup 1, when all CT findings were used for CT–based diagnosis of TMJ OA, 65 joints were identified as affected. Among these, TMJ OA was confirmed histopathologically in only 63% of joints (41/65 TMJs). Radiological assessment based on all CT findings resulted in high sensitivity (1.0) but low specificity (0.41), indicating that 24 TMJs were falsely diagnosed as OA–affected.

**Table 6.** The accuracy of CT–based recognition of temporomandibular joint osteoarthritis (TMJ OA) in relation to joints with histopathologically confirmed TMJ OA. Three subgroups are considered: subgroup 1 – including all CT findings; subgroup 2 – including CT findings excluding ‘CT anatomical variations’ [20], and subgroup 3 – including CT findings with bone cysts [28] but excluding other ‘CT anatomical variations’ [20].

| Accuracy metrics | TMJ OA-related groups |  |  |
| --- | --- | --- | --- |
|  | Subgroup 1 | Subgroup 2 | Subgroup 3 |
| Recognized / Confirmed; n / n (%) | 65 / 41 (63%) | 26 / 23 (88%) | 35 / 32 (91%) |
| Se | 1.00 | 0.56 | 0.79 |
| 95% CI | 0.91 to 1.00 | 0.40 to 0.72 | 0.62 to 0.89 |
| Sp | 0.41 | 0.93 | 0.92 |
| 95% CI | 0.26 to 0.58 | 0.80 to 0.98 | 0.80 to 0.98 |
| AUC | 0.71 | 0.74 | 0.85 |
| 95% CI | 0.60 to 0.80 | 0.64 to 0.83 | 0.76 to 0.92 |
| PPV | 0.63 | 0.89 | 0.91 |
| 95% CI | 0.57 to 0.69 | 0.71 to 0.96 | 0.78 to 0.96 |
| NPV | 1.00 | 0.68 | 0.81 |
| 95% CI | 0.80 to 1.00 | 0.60 to 0.75 | 0.70 to 0.88 |

Footnotes: AUC = area under curve; NPV = negative predictive value; PPV = positive predictive value; Se = sensitivity; Sp = specificity; n / n = number of TMJ with at least one CT finding (recognized) in relation to number of true positive TMJ OA (confirmed); % = percent of histopathologically confirmed TMJ OA among CT-based recognized TMJ OA; 95% CI = 95% confidence interval

In subgroup 2, when CT findings excluding ‘CT anatomical variations’ [20] were used for CT–based diagnosis of TMJ OA, 26 joints were identified as affected. Among these, TMJ OA was confirmed histopathologically in 88% of joints (23/26 TMJs). This radiological assessment resulted in high specificity (0.93) but low sensitivity (0.56), indicating that 15 TMJs were falsely diagnosed as OA–free.

In subgroup 3, when CT findings including bone cysts [28] but excluding other ‘CT anatomical variations’ [20] were used for CT–based diagnosis of TMJ OA, 35 joints were identified as affected. Among these, TMJ OA was confirmed histopathologically in 91% of joints (32/35 TMJs). This radiological assessment resulted in moderate sensitivity (0.79) and high specificity (0.92), indicating that only 3 TMJs were falsely diagnosed as OA–affected, while 9 TMJs were falsely diagnosed as OA–free.

## 4 DISCUSSION

CT imaging of the equine head is successfully used in clinical practice to support the diagnosis of TMJ OA [3]. Numerous CT findings have recently been reported both in horses with TMJ OA [5–7,14,15] and incidentally in asymptomatic horses [20], raising the question: ‘When do CT findings become evidence of TMJ disease in an apparently clinically normal horse?’ [20].

To date, only five case reports have simultaneously described radiologic and histopathological findings in individual horses, including two reports of TMJ OA [5,14], one report of septic TMJ arthritis [36], one of squamous cell carcinoma [37], and one of a dentigerous cyst located in the TMJ region [38]. Moreover, among the research studies that have histopathologically examined equine TMJ tissues [32,39– 42], only one specifically addressed TMJ OA, although it did not incorporate CT imaging [39]. The remaining four studies described normal articular cartilage [32,40–42] and the intra–articular disc [39], while only studied histopathological findings jointly with micro–CT (μCT) images [41,42]. Several additional case reports [6,7] and research studies [15,18,20,30] have investigated CT imaging of both normal [20,30] and OA–affected [6,7,15,18] TMJs; however, none included histopathological analysis. Therefore, to the best of our knowledge, this is the first prospective study to investigate the relationship between CT findings and histopathological manifestations of OA in equine TMJs.

In the first of these two case reports, Pimentel et al. [14] described unilateral TMJ dysplasia with osteochondrosis–like lesions and associated OA in a 15–month–old filly. CT findings included flattening of the mandibular condyle, osseous fragments within the joint space, medial and rostral osteophytes, and mineralization of the intra–articular disc [14]. Three of these CT findings – flattening, medial osteophytes (which should be termed medial enthesophytes [20]), and disc mineralization – were suggested as ‘CT anatomical variations’ [20], while the fourth – osseous fragments within the joint space – may represent a manifestation of the osteochondrosis observed in this filly [14]. In this case report, histopathology revealed foci of necrosis, chondrocyte hypertrophy, endochondral ossification, multiple cyst–like defects, and fibrosis in the articular cartilage of the mandibular condyle [14], among which only chondrocyte necrosis was listed in the OARSI recommendations for histological assessment of OA in horses [31].

In the second case reports, Smyth et al. [5] described an 18–year–old mare with unilateral CT findings and bilateral histopathological evidence of TMJ OA. CT findings included subchondral sclerosis of the mandibular condyle, medial osteophytes of the mandibular condyles (which should be termed medial enthesophytes [20]), lateral osteophytes of the mandibular fossa, and mineralization of the intra–articular disc [5]. Three of two CT findings – medial entesiophytosis and disc mineralization – were suggested as ‘CT anatomical variations’ [20]. In this case report, histopathological analysis revealed cartilage degeneration and chondrones in both the mandibular condyle and mandibular fossa articular cartilage, as well as chondroid and chondro–osseous metaplasia in the intra–articular disc [5], among which only chondrones (cluster formation) were listed in the OARSI recommendations [31]. Since both publications were single case reports [5,14], assessing the relationship between CT findings and the histopathological manifestations of OA has not been possible so far.

Among the CT findings described in the case reports by Pimentel et al. [14] and Smyth et al. [5], subchondral sclerosis and intra–articular disc mineralization were not observed in the present study. Osteophytes were identified on the lateral margin of the mandibular condyle in the present study, whereas Smyth et al. [5] reported them on the lateral margin of the mandibular fossa and Pimentel et al. [14] on the rostral margin of the mandibular condyle. Lateral osteophytes showed only a weak positive correlation with subintimal edema in the joint capsule and no significant association with the horses’ age. Furthermore, flattening and medial enthesophytes of the mandibular condyle were positively correlated with chondrocyte necrosis, cluster formation, and partial focal cell loss in the articular cartilage, as well as with numerous alterations in the joint capsule. These CT findings were observed only in horses older than 5 years, similar to the adult mare described by Smyth et al. [5], but contrary to the filly reported by Pimentel et al. [14]. In addition, these findings showed strong positive correlations with age, supporting the notion that primary OA in horses may be associated with progressive wear–and–tear of the articular cartilage over time [5,6,20,27]. Histopathological manifestations of wear–and–tear of the articular cartilage have also been described in clinically normal horses of various ages [32]. Horses younger than 10 years exhibited fewer histopathological alterations in both the articular cartilages and intra–articular disc compared with horses older than 20 years; however, specific histopathological findings were not analyzed [32]. Age–related differences in the appearance of TMJ articular cartilage, as summarized by Smyth et al. [32], were well illustrated by Mirahmadi et al. [41]. Mirahmadi et al. [41] demonstrated that, with increasing age, the articular cartilage of the mandibular condyle becomes thinner, whereas collagen, glycosaminoglycan, and pentosidine content increase and water content decreases. However, neither study [32,41] described age– related change in the individual histopathological findings listed in the OARSI recommendations [31].

In the present study, chondrocyte necrosis and cluster formation were positively correlated with the horses’ age in the articular cartilage of both the mandibular condyle and mandibular fossa. It should be noted, however, that unlike appendicular joints – where the articular surface is covered by hyaline cartilage [43] – the TMJ is lined with fibrocartilage [40]. Consequently, structural changes in the aging articular cartilage of the mandibular condyle may differ from those observed in hyaline cartilage [41]. In the equine TMJ, the articular surfaces are directly underlain by a zone of fibrocyte–like cells [40], which is absent in appendicular joints [6]. This zone has been proposed to enable continuous fibrocartilage regeneration [6,20], similar to that described in humans [44]. However, TMJ fibrocartilage is not capable of complete regeneration; therefore, articular cartilage alterations accumulate over time and become more frequent in older horses [32]. The present results provide evidence that the equine TMJ undergoes age–related remodeling, the sings of which are correlated and visible on both histopathological [32,41] and CT [20] images, although these findings have been reported independently in previous studies [20,32,41].

It should be noted that certain CT findings, particularly those classified as ‘CT anatomical variations’ [20], were associated with age, as reflected by differences in their frequency distributions and age–related correlations. For example, scattered hypodense regions in the mandibular condyle were observed only in horses younger than 5 years and showed a moderate negative correlation with horse age, as well as weak negative correlations with two alternations in the articular cartilage and three alterations in the joint capsule.

Moreover, the observed negative correlations suggest that this CT finding may be consistent with physiological remodeling of the mandibular condyles in young horses and may occur in asymptomatic individuals as part of normal mineralization [20]. Notable, scattered hypodense regions have not been described in other studies of symptomatic TMJ dysfunction [3], supporting the assumption that this finding is not indicative of OA. These observations may suggest a link between mineralization of the mandibular condyle in young horses and the presence of scattered hypodense regions, potentially representing a marker of immaturity, since changes in the mandibular condyles of skeletally immature horses may reflect ongoing maturation [20]. Therefore, practitioners performing CT examinations in young horses should interpret CT findings with caution [20], particularly when septic arthritis [1,10,17] or dysplasia with osteochondrosis–like lesions [14] are clinically suspected.

In contrast, flattening and medial enthesophytosis of the mandibular condyle – also categorized as ‘CT anatomical variations’ [20] – showed strong positive correlations with age. Moreover, these CT findings demonstrated weak to strong positive correlations with alterations in both the articular cartilage and the joint capsule, which may reflect age–related wear–and–tear [20,32]. Interestingly, these two CT findings have also been reported in horses with clinical symptoms of TMJ dysfunction; however, they were always accompanied by other CT findings considered indicative of OA in humans [28] – such as irregularity of the bone surface [15], lateral [5] or rostral [14] osteophytosis, subchondral sclerosis [5], or subchondral bone cysts [6,7,15]. Therefore, flattening and medial enthesophytosis of the mandibular condyle, in the absence of more specific CT findings, likely represents an age–related phenomenon [20]. This supports the conclusion that not all abnormalities observed on CT images of the equine head are indicative of TMJ OA.

It should be emphasized that ‘CT anatomical variations’ of the mandibular condyle – particularly flattening, scattered regions of hypodensity, subchondral bone cyst without surrounding sclerosis, and medial enthesophytes – were observed in TMJs both with and without histopathologically confirmed OA. Consequently, when all CT findings were included in the CT–based diagnosis, sensitivity was low. Inclusion of all CT findings resulted in a high rate of false–positive diagnoses, leading to overdiagnosis of TMJ OA on CT. This was likely due to the inclusion of CT findings associated which normal mineralization of the mandibular condyle in young horses and age–related remodeling of the mandibular condyle in older horses [20]. In contrast, exclusion of all ‘CT anatomical variations’ resulted in a substantial number of false– negative diagnoses, leading to underdiagnosis of TMJ OA. However, when subchondral bone cysts with surrounding sclerosis were included in accordance with the DC/TMD guidelines [28], sensitivity reached a moderate value of 0.79 with maintaining high specificity.

Although subchondral bone cysts have been described as incidental findings in asymptomatic horses [20], they have also been reported as clinically significant [6,7,15] and have been surgically treated with long–term follow–up [6,15]. Notingworth, subchondral bone cysts in appendicular joints are associated with OA and its dysfunction manifesting as horse’s lameness [45]. Also in the mandibular condyle, subchondral bone cyst has been reported in a horse exhibiting signs of TMJ dysfunctions, such as ‘clunking’ during mastication [6], as well as signs of mild TMJ–related pain, such as aversive behavior [6] and poor performance [7]. Accordingly, Carmalt and Reisbig (2022) noted that TMJ bone cysts become clinically relevant when associated with synovitis and OA [6], resulting in clinically detectable TMJ–related pain [6,7]. The authors further suggested that subchondral bone cysts and TMJ OA may share a similar etiology: the former may arise from trauma to the subchondral bone plate or developmental anomalies, whereas the latter may result also from trauma as well as septic arthritis or chronic wear–and–tear [6]. Although the relationship between TMJ bone cysts, OA, and clinical symptoms remains unclear [6,20], including bone cysts with surrounding sclerosis in CT–based diagnostic criteria appears to improve diagnostic accuracy of TMJ OA compared with assessments based solely on irregularity of the mandibular condyle, lateral osteophytosis, and joint space narrowing. However, the clinical significance of these CT findings requires further investigation.

One limitation of this study is that gross examination of the TMJs was not performed, although this should be included in future studies in accordance with the categories described by McIlwraith [31]. Among the studies that have histopathologically examined equine TMJ tissues [32,39–42], only one included a gross examination of the TMJ [39], which served as the basis for classifying TMJs as normal, mild OA, or severe OA. However, this subjective classification did not account for the OARSI recommendations [31]. A second limitation is that the effect of gender on CT and histopathological findings was not evaluated in this study due to the small sample size. It should be noted that in a previous study, asymptomatic male horses were more likely to have bone cysts in the mandibular condyle than females, while gender had no effect on the presence of intra–articular disc mineralization [20]. However, these findings were based on CT scans of 1018 horses [20], whereas in the present study, only 50 cadaver heads were examined, of which only 41 underwent the full experimental protocol. Therefore, future studies designed with larger sample sizes of cadaver heads should also consider the gender effect. A third limitation is the absence of clinical data for the horses whose heads were examined. Because the heads were collected post–mortem from a slaughterhouse, clinical signs could not be assessed. However, even in the retrospective clinical study involving a large cohort of asymptomatic horses, follow–up data on behavior during eating or riding were unavailable. Therefore, the presence of mild TMJ–related pain cannot be definitively excluded. The authors suggested that some horses with ‘CT anatomical variations’ may have had subclinical TMJ disease without exhibiting moderate or severe clinical signs of pain, especially as 10.7% of the studied horses underwent CT examination due to neurological problems, including head–shaking syndrome [20]. Given that, in other studies, horses with head–shaking syndrome were suspected to have TMJ OA [5,18], the clinical significance of specific CT findings in equine TMJs warrants further investigation.

## 5 CONCLUSIONS

Given that equine TMJ underwent age–related remodeling, not all findings observed on CT images of the horse’s head are indicative of TMJ OA. Some CT findings in TMJ, particularly those referred to as ‘CT anatomical variations’, showed an association with age, as evidenced by differences in their frequency distributions and age–related correlations. As ‘CT anatomical variations’ – particularly flattening of the mandibular condyle, scattered regions of hypodensity in the mandibular condyle, subchondral bone cyst without surrounding sclerosis, and medial enthesophytes in the mandibular condyle – were observed in TMJs with histopathologically confirmed OA and OA–free, inclusion of all CT findings in the CT–based diagnosis of TMJ OA yielded a high rate of false–positive diagnoses. In contrast, exclusion of all ‘CT anatomical variations’ resulted in a substantial number of false–negative diagnoses. Therefore, it may be suggested that inclusion of subchondral bone cysts with surrounding sclerosis – classified among ‘CT anatomical variations’ – into CT–based diagnosis of TMJ OA increases its accuracy compared to TMJ assessment considering only irregularity of the mandibular condyle, lateral osteophytosis, and narrowing of the joint space. However, the clinical significance of these results remains unknown and requires further research.

